# Denitrification genotypes of endospore-forming *Bacillota*

**DOI:** 10.1101/2024.05.17.594689

**Authors:** Emma Bell, Jianwei Chen, Milovan Fustic, Casey RJ Hubert

## Abstract

Denitrification is a key metabolic process in the global nitrogen cycle and is performed by taxonomically diverse microorganisms. Despite the widespread importance of this metabolism, challenges remain in identifying denitrifying populations and predicting their metabolic end-products based on their genotype. Here, genome-resolved metagenomics was used to explore the denitrification genotype of *Bacillota* enriched in nitrate-amended high temperature incubations with confirmed N_2_O and N_2_ production. A set of 12 hidden Markov models (HMMs) was created to target the diversity of denitrification genes in members of the phylum *Bacillota*. Genomic potential for complete denitrification was found in five metagenome-assembled genomes (MAGs) from nitrate-amended enrichments, including two novel members of the *Brevibacillaceae* family. Genomes of complete denitrifiers encode N_2_O reductase *(nos)* gene clusters with clade II-type *nosZ* and often include multiple variants of the nitric oxide reductase *(nor)* gene. The HMM set applied to all genomes of *Bacillota* from the Genome Taxonomy Database (GTDB) identified 17 genera inferred to contain complete denitrifiers based on their gene content. Among complete denitrifiers it was common for three distinct nitric oxide reductases to be present (qNOR, bNOR and sNOR) which may reflect the metabolic adaptability of *Bacillota* in environments with variable redox conditions.

## Introduction

Denitrification is a key metabolic process in the nitrogen cycle featuring sequential reduction of nitrate to nitrite and then gaseous metabolites (NO_3_^-^ → NO ^-^ → NO → N_2_O → N_2_). Different enzymes catalyse each of the four reduction reactions such that this modular metabolism can be performed by a single microorganism or a microbial consortium. When performed modularly, microorganisms can achieve complete denitrification by cross-feeding intermediates. If denitrification is incomplete, this can give rise to the release of the greenhouse gas N_2_O to the atmosphere [1, 2]. Denitrification is prevalent in terrestrial and aquatic environments where oxic and anoxic conditions occur close to each other [3]. In soil environments, denitrification can contribute to losses of fixed nitrogen to the atmosphere reducing soil fertility and plant yield [4]. On the other hand, denitrification in wetlands mitigates the transport of nitrogen from land to lakes and coastal waters where excess nitrogen can cause eutrophication [5]. Nitrogen removal in wastewater and agricultural sectors via denitrification similarly represents a critical step that limits the release of excess to nitrogen into watersheds [6, 7]. Identifying microorganisms contributing towards denitrification in different environments is therefore of ecological and industrial importance.

Microorganisms known to perform denitrification are taxonomically diverse and span both bacterial and archaeal domains [8, 9]. The taxonomic diversity of denitrifiers means that this metabolism cannot be easily linked to phylogeny [10]. Instead, characterisation of denitrifying populations relies on the presence of metabolic genes for each of the reduction steps. Nitrate reduction to nitrite (NO_3_^-^ → NO_2_^-^) is catalysed by the membrane-bound Nar enzyme or periplasmic Nap enzyme, both of which can be found in denitrifiers and nitrate-ammonifiers (NO_3_ ^-^ → NH_4_ ^+^). Thus, the presence of nitrite reductase (Nir) genes for nitrite reduction to nitric oxide (NO ^-^ → NO), nitric oxide reductase (Nor) genes for nitric oxide reduction to nitrous oxide (NO → N_2_O), and the nitrous oxide reductase (Nos) gene for nitrous oxide reduction to dinitrogen (N_2_O → N_2_) are used as markers that differentiate denitrification from dissimilatory nitrate reduction to ammonium (DNRA).

Enzymes from the denitrification pathway exhibit broad taxonomic and sequence diversity. Nitrite reduction to nitric oxide is catalysed by two structurally different enzymes, Cu-type NirK and cytochrome cd-1 type NirS, that have different evolutionary histories [11–13]. Nitric oxide reductases are members of the heme-copper oxidase (HCO) superfamily and are ancestral to terminal oxidases for aerobic respiration [14, 15]. Four nitric oxide reductase enzyme families have been biochemically characterised: cNOR [16], qNOR [17], bNOR (formerly Cu_A_NOR; [18]) and eNOR [19]. Cytochrome *c*-dependent (cNOR) and quinol-dependent (qNOR) Nor enzymes are related to C-family oxygen reductases whereas bNOR and the recently characterised eNOR are related to B-family oxygen reductases [15, 19]. Based on phylogenomic analysis and conserved proton channels, an additional three enzymes related to B-family oxygen reductases, sNOR, gNOR, and, nNOR, have also been proposed [19].

In contrast to nitric oxide reduction, nitrous oxide reduction is catalysed by just one enzyme, NosZ. This enzyme, however, forms two distinct groups known as clade I (typical) or clade II (atypical) characterised by different secretionary pathways (tat and sec, respectively) [9, 20]. Clade I is represented by well-studied denitrifying *Proteobacteria* (alpha-, beta-, and gamma-) whereas clade II is taxonomically diverse and encompasses at least 12 bacterial and archaeal phyla (subclades A−K) [9, 21].

The broad diversity and sequence divergence among denitrification enzymes gives rise to well-documented coverage limitations for PCR primers and probes [9, 10, 12, 22, 23]. This makes it challenging to accurately estimate the diversity and abundance of denitrification genes. Metagenomics circumvents primer bias limitations and is therefore advantageous for studying denitrification. Here, genome-resolved metagenomics was employed to explore the gene content of microorganisms enriched in the presence of nitrate from oil sands outcrops in Alberta, Canada. Compared to conventional crude oil reservoir ecosystems, oil sands are not well characterised microbiologically but the presence of both mesophilic and thermophilic populations in riverbank outcrops and subsurface deposits has been reported [24, 25]. Our data show that dormant thermophilic endospore-forming (thermospore) populations with distinct denitrification genotypes are present in the oil sands microbiome.

## Methods

### Sample collection

Samples were collected from the Athabasca oil sands in Alberta, Canada, in June 2019. In this region, oil sands are present at various depths (up to 100s of metres) and outcrops are naturally exposed along the riverbanks of the Athabasca River and its tributaries. Oil sands samples were collected from an outcrop at the Hangingstone River (56°42’38"N, 111°23’51") in Fort McMurray. Samples were stored in a cold room at 4°C until incubations were established. Parallel samples were frozen for DNA extraction and analysed to represent unincubated samples.

### Nitrate-amended enrichments

Approximately 30 g oil sands inoculum was combined with 60 mL anoxic medium in 160 mL Wheaton glass serum bottles. Growth medium was based upon media previously used to isolate thermophilic *Bacillota* (formerly *Firmicutes*) from hydrocarbon environments (Adkins *et al.*, 1992; Salinas *et al.*, 2004) and contained (L^-1^ distilled water): 0.2 g MgCl_2_•6H_2_O, 0.1 g KCl, 1 g NH_4_Cl, 0.1 g CaCl_2_•2H_2_O, 0.3 g K_2_HPO_4_, 0.3 g KH_2_PO_4_, 1 g NaCl, 0.2 g yeast extract. NaNO_3_ (24 mM) was added as an electron acceptor and glucose (5 mM) was added as an electron donor. Cysteine hydrochloride (2.8 mM), NaHCO_3_ (0.3 M), vitamins and trace minerals were added from sterile stock solutions. Anoxia was established with He to enable sub-samples of headspace gas to be analysed for the presence of nitrogen compounds. Throughout the incubation period, sub-samples of the sand and water mixture (1.5 mL) were periodically removed from enrichments using a He-flushed syringe. Sub-samples were centrifuged (10,000 × g for 5 minutes) with supernatants filtered (0.2 µm) and frozen eventual for chemical analysis and pellets frozen for eventual DNA extraction.

### Headspace gas measurement

Headspace gases (1 mL) were extracted from experimental incubations with a He-flushed syringe and immediately injected into two chain-connected sample loops on an Agilent 7890B gas chromatograph (GC). CO_2_ was first separated on a Hayesep N packing column (stainless steel tubing, 0.5 m length, 1/8″ OD, 2 mm ID, mesh size 80/100) followed by N_2_ separation on a MolSieve 5A packing column (UltiMetal tubing, 2.44 m length, 1/8″ OD, 2 mm ID, mesh size 60/80) with He carrier gas. Both CO_2_ and N_2_ were measured by thermal conductivity detector (TCD) at 200°C. Through a second line, N_2_O was separated on a Hayseed Q packing column (stainless steel tubing, 6’ length, 1/8″ OD, 2.1 mm ID, mesh size 80/200) with Ar/CH_4_ 5/95% carrier gas. N_2_O was measured by electron capture detector (ECD) at 300°C. All columns were set in the same oven with a working temperature of 105°C.

### Chemical analyses

Nitrate and nitrite were measured with a Dionex ICS-5000 reagent-free ion chromatography system equipped with an anion-exchange column (Dionex IonPac AS22; 4 x 250 mm). The eluent was 4.5 mM K_2_CO_3_ / 1.4 mM KHCO_3_, the flow rate was 1.3 mL/min, and the column temperature was 30°C. Organic acids (formate, acetate, propionate, lactate, butyrate and succinate) were measured using UV (210 nm) on an HPLC RSLC Ultimate 3000 equipped with an Aminex HPX-87H, 7.8 x 300 mm analytical column. The isocratic eluent was 5 mM H_2_SO_4_, the flow rate was 0.6 mL/min, and the column oven was heated to 60°C. Glucose was measured by mixing samples with the Glucose (HK) Assay reagent (Sigma-Aldrich) following the manufacturer’s instructions and absorbance was measured on a spectrophotometer at 410 nm.

### DNA extraction

DNA was extracted from frozen pellets (0.25 g) using the Qiagen DNeasy PowerLyzer PowerSoil kit according to the manufacturer’s protocol. DNA concentrations were measured using the dsDNA High Sensitivity assay kit on a Qubit 2.0 fluorometer. DNA yields from heated oil sands ranged between 329– 7890 ng DNA g^-1^ oil sand. To represent unincubated samples, DNA was extracted 8-10g oil sands (i.e., samples that were frozen when original outcrop samples were collected) using the Qiagen DNeasy PowerMax Soil kit according to the manufacturer’s protocol. Triplicate DNA extractions from these oil sands yielded 172–228 ng DNA g^-1^.

### 16S rRNA gene amplicon sequencing

Amplicon sequencing of the 16S rRNA gene (V4−V5 region) was performed using the bacterial primer set 515F and 926R [26]. Triplicate PCR reactions were pooled then purified using the NucleoMag NGS clean-up and size select kit. Purified PCR products were indexed following Illumina’s 16S rRNA amplicon preparation instructions. Indexed amplicons were verified on an Agilent 2100 Bioanalyzer system and sequenced on a MiSeq benchtop sequencer (Illumina) using the v3 600-cycle (paired end) reagent kit. Primers were trimmed from sequence reads with Cutadapt v4.4 [27] and processed in DADA2 [28] following the recommended pipeline (https://benjjneb.github.io/dada2/tutorial.html). Taxonomy was assigned with ‘assignTaxonomy’ in DADA2 using the Swedish Biodiversity Infrastructure (SBDI) Sativa curated 16S GTDB database from release R07-RS207 (https://doi.org/10.17044/scilifelab.14869077).

### Metagenome sequencing, read processing, and binning

Metagenomic sequencing was performed on a NovaSeq 6000 (Illumina) with a S4 300 cycle flow cell. Libraries were prepared by shearing to an insert size of 200 bp using a Covaris instrument followed by library construction with the NEB Ultra II DNA library prep kit. Adaptors and low-quality reads were removed with Cutadapt v.1.18 [27] using the wrapper Trimgalore v0.6.7 [29]. Reads from each sample were assembled individually with Megahit v1.2.9 [30] using the ‘--meta-sensitive’ option. Read and assembly statistics are provided in **Table S1**. Reads were cross-mapped to the assembled contigs with BBMap v38.95 [31] to generate coverage profiles for binning. Contigs from each assembly were then binned with MetaBAT2 [32] and CONCOCT [33] and refined with DAS Tool [34]. Bin completeness and contamination were calculated with CheckM2 [35]. Bins with >50% completeness and <10% contamination were dereplicated with dRep [36] resulting in 17 non-redundant metagenome assembled genomes (MAGs) (**Table S2**). Relative abundance of non-redundant MAGs was determined with CoverM using default parameters for ‘coverm genome’ [37].

### Annotation and evaluation MAGs

MAGs were taxonomically classified with Genome Taxonomy Database Toolkit (GTDB-tk) v2.3.2 with reference data R214 [38]. Average amino acid identity (AAI) comparisons between MAGs without close relatives in GTDB (<70% AAI) were determined with AAI calculator [39]. Functional annotation with KEGG and EggNOG databases was performed with GhostKOALA [40] and eggnog-mapper v2.1.7 [41]. Optimal growth temperature was predicted with from the genome sequences with ‘tome predOGT’ [42] (**Table S2**) and MAGs were checked for functional and regulatory genes involved in endospore formation (**Table S3**). MAGs of mesophilic non-endosporulating bacteria included *Actinobacteriota* (×1), *Patescibacteria* (×2) and *Proteobacteria* (×2). These MAGs binned from unheated oil sands inoculum and had low relative abundance in heated samples so were excluded from further analysis. The final non-redundant set of thermospore MAGs contained 12 high quality genomes (**Table S2**).

### Annotation of the Nos gene cluster

Genes identified as *nosZ* were checked for the presence of Sec/SPI signal peptides, characteristic of all clade II *nosZ*, with SignalP 6.0 [43]. Genes on the same contig as *nosZ* were then checked for transmembrane helices with DeepTMHMM [44] to identify NosB which is typically found adjacent to NosZ in genomes with clade II *nosZ* and contains 4 or 6 transmembrane helices [45]. A cytochrome *c* was identified preceding *nosZ* with eggNOG and was determined *nosC* [46]. Cellular localisation of denitrification genes was predicted with PSORTb. [47]. Nos gene clusters were visualised with the gggenes extension [48] for gglot2 [49].

### Hidden Markov models (HMMs) for denitrification

A denitrification gene set was compiled from both custom HMMs and TIGRfams (**Table S4**). Six HMMs were created for *nirK*, *nirS*, *qnor*, *bnor*, *snor* and *nosB.* Nucleotide sequences from genes of interest were retrieved from studies biochemically characterising and describing the enzymes [12, 17, 19, 50, 51]. Multiple sequence alignments of characterised nucleotide sequences and related nucleotide sequences from MAGs in this study were created with Clustal-Omega v1.2.4 [52]. Each alignment was manually inspected for conserved active site residues essential for structure and function in AliView v1.28 [53]. HMMs were created from the inspected multiple sequence alignments with ‘hmmbuild’ in HMMER 3.3.2 [54]. The HMMs were first tested on MAGs from this study using ‘hmmsearch’ and the trusted cutoff (TC) values were iteratively adjusted to ensure only genes with conserved residues were captured. The resulting HMMs were used to retrieve denitrification gene sequences from *Firmicutes* genomes (*Firmicutes* and *Firmicutes* A−H, *n* = 13,543) downloaded from GTDB R07-RS207 [55] using ‘gtt-get-accessions-from-GTDB’ in GToTree v1.6.34 [56]. Note that *Firmicutes* phyla were renamed *Bacillota* following the release of GTDB reference data R214. Denitrification gene sequences from GTDB and thermospore MAGs were dereplicated with ‘fastx_uniques’ in USEARCH v11 [57]. Unique sequences were aligned with Clustal-Omega v1.2.4 [52], manually inspected, and included in a revised HMM. TIGRfams were used to identify the genes *narGH*, *napA*, *nosZI* (clade I) and *nosZII* (clade II) [58]. Following manual inspection, the trusted cutoff for the *napA* HMM was amended to capture monomeric NapA found in *Bacillota E* genomes. The final HMM set is available at https://github.com/emma-bell/metabolism.

### Visualisation of denitrification genes in *Bacillota*

*Bacillota* genomes from GTDB with Nor and/or Nos genes identified in their genome (*n* = 433) were visualised in a phylogenomic tree with thermospore MAGs from this study. The tree was created with GToTree v1.6.34 [56] from a concatenated alignment of 119 single copy genes targeted by the *Firmicutes* HMM profile. Genes were first predicted with Prodigal v2.6.3 [59] and target genes were identified with HMMER3 v3.3.2 [54]. Target genes were individually aligned with muscle v5.1 [60] and trimmed with TrimAl v1.4.rev15 [61]. Concatenated sequence alignments were used to create the phylogenomic tree with FastTree2 v2.1.11 [62]. The tree was transformed and annotated in Treeviewer v2.2.0 [63].

## Results

### Enrichment of thermophilic denitrifiers

Incubation of oil sands at 50°C with nitrate and glucose resulted in nitrogen compounds being sequentially reduced (**Fig. 1A–C**) coupled to the metabolism of glucose into organic acids and carbon dioxide (**Fig. 1D–F**). N_2_ gas production and organic acid consumption was variable between replicates, with 16S rRNA gene amplicon sequencing confirming that distinct populations were enriched in different bottles (**Fig. 1 G–I**). Amplicon sequence variants (ASVs) from the genera *Brevibacillus*, *Neobacillus*, *Geobacillus*, *Paenibacillus* and JAGHKQ01 (family *Bacillaceae G*) were found in common across triplicate enrichments, whereas other taxonomic lineages were exclusive. For example, ASVs of the class *Bacilli* (ASVs 6 and 18) were enriched only in incubation 1 (**Fig. 1G**) whereas ASVs of the family *Brevibacillaceae* (ASVs 9, 33, 35) and genus *Symbiobacterium* (ASVs 4, 15, 24, 25, 27) were enriched only in incubation 2 (**Fig. 2H**). This experimental approach therefore showed potential to uncover a diverse range of thermophilic denitrifiers, with different members of the oil sands microbial seed bank becoming enriched from within parallel inocula.

**Figure 1:**
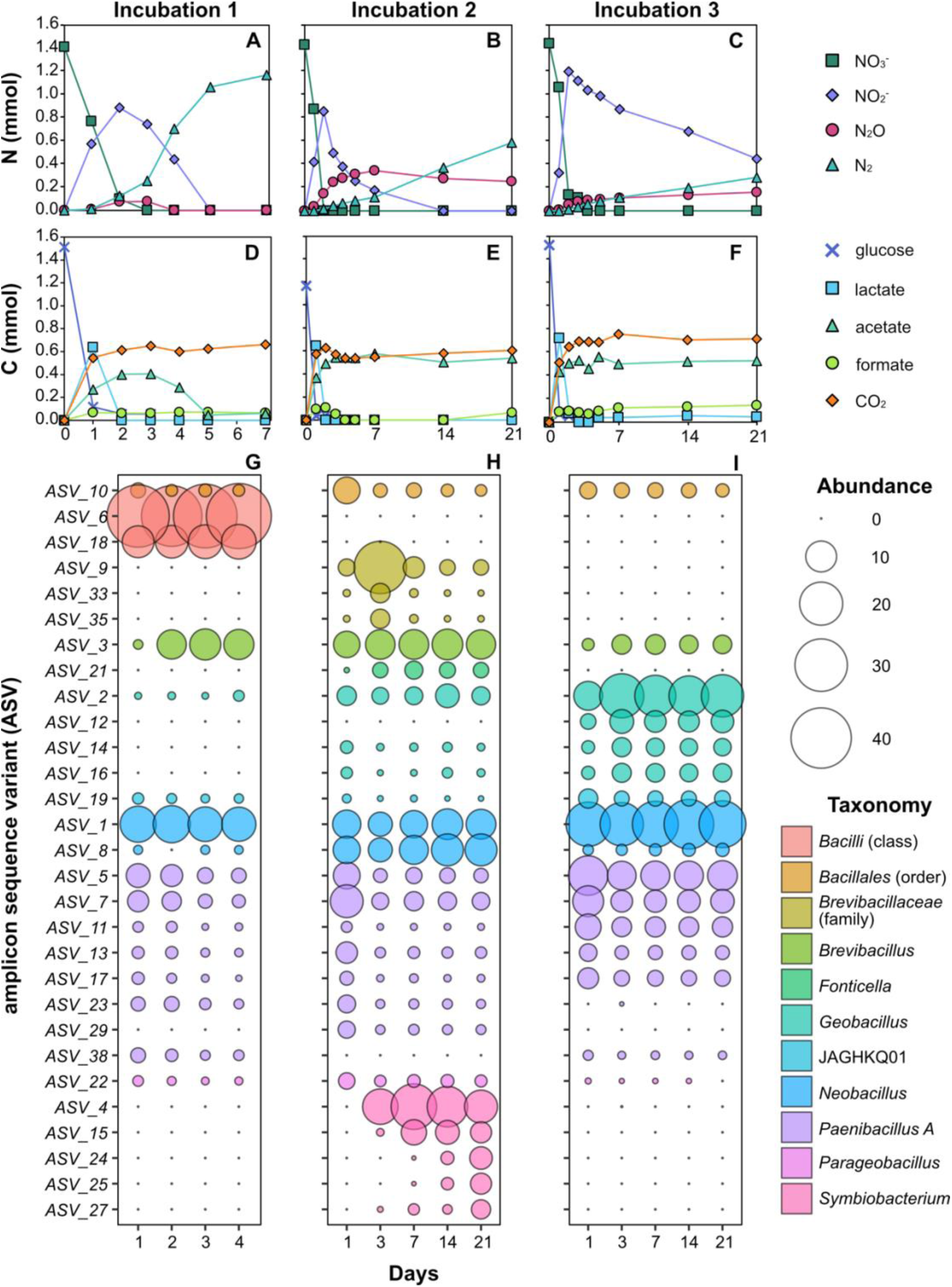
Thermospores enriched in heated nitrate-amended incubations. Nitrate reduction and production of nitrite, nitrous oxide and dinitrogen was monitored during incubation at 50°C (**A**, **B**, **C**) (nitric oxide was not measured). Glucose, organic acids, and CO_2_ measurements for each incubation are shown in corresponding panels underneath (**D**, **E**, **F**). 16S rRNA gene amplicons were sequenced from multiple time points (**G**, **H**, **I**) and *Bacillota* represented 94–98% read abundance in all cases. For clarity, only amplicon sequence variants (ASVs) detected at >2% read abundance are included in the plots.

**Figure 2:**
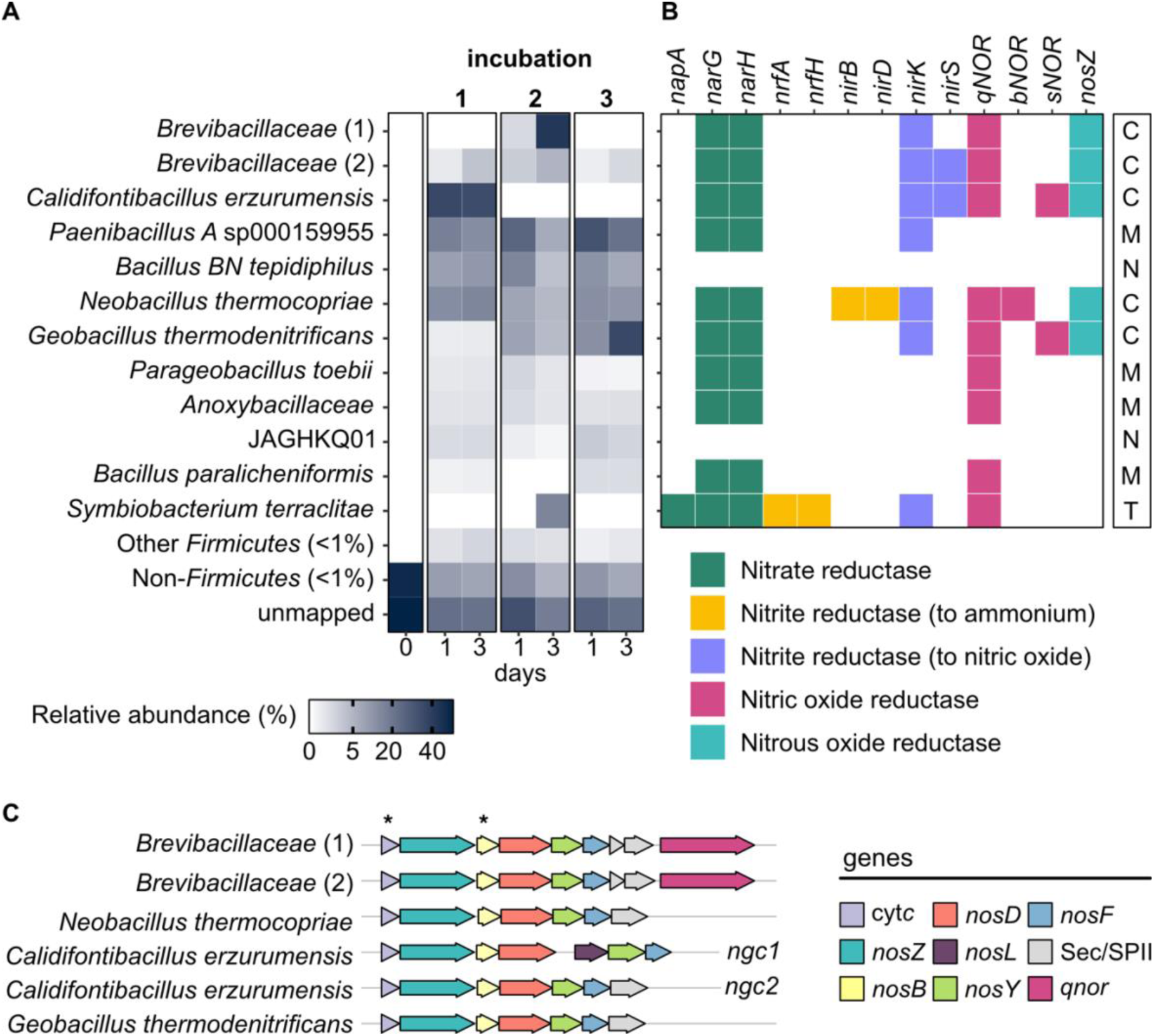
Denitrification genotypes of enriched thermospores. Relative abundance of thermospore MAGs in incubations with NO_3_^-^ after one- and three-days incubation at 50°C **(A)**. For comparison, the abundance of thermospores in the inoculum, prior to their enrichment, is shown as 0 days. Based on denitrification genes present **(B)**, MAGs were classified as complete (C), truncated (T), modular (M) or non-denitrifying (N) with respect to their potential for denitrification (column at right). Complete denitrifiers encode clade II-type NosZ. The Nos gene cluster (ngc) of thermospore MAGs is shown **(C)** including two gene clusters in the single *Calidifontibacillus erzurumensis* MAG. Asterisks indicate conserved genes found in clade II.

### Reconstruction of thermospore metagenome-assembled genomes (MAGs)

Metagenomic sequencing was performed on the inoculum (i.e., unheated oil sands) and on the sub-samples taken from the 50°C incubations after 1 and 3 days. Twelve high quality MAGs from *Bacillota* phyla (*Bacillota* ×11, *Bacillota E* ×1) were recovered from heated incubations. Reads from the *Bacillota* MAGs were not detected in the unheated oil sands metagenome (**Fig. 2A**) which is consistent with dormant thermospore populations only germinating upon heating. Genomic potential for endospore formation and germination in *Bacillota* MAGs was also confirmed by the presence of functional and regulatory genes conserved within endospore-forming taxa (**Table S2**).

### Identifying genes for denitrification

KEGG orthologs (KO) did not capture the diversity of denitrification enzymes present in *Bacillota*. With KEGG, the nitric oxide reductase qNOR was annotated as subunit B of cNOR (*norB*, K04561) and nitric oxide reductases bNOR and sNOR are annotated as the related but functionally distinct cytochrome *c* oxidase (*coxA*, K02274). Nitrous oxide reductase is identified with KEGG (*nosZ*, K00376) but clade I and clade II enzymes are not differentiated. Furthermore, the only gene annotated in addition to *nosZ* from the Nos gene cluster was the accessory protein *nosD* (K07218), found in both clade I and clade II *nosZ* microorganisms [64]. A set of HMMs was therefore created to capture denitrification potential in genomes of *Bacillota*. The HMM set includes distinct HMMs for nitric oxide reductases and differentiates between clade I and clade II NosZ. An HMM for the membrane lipoprotein *nosB* is also included. NosB is essential for N_2_O respiration in clade II *nosZ* microorganisms but is commonly absent in clade I microorganisms [21, 45].

Using the HMM set, MAGs were designated as complete-, truncated-, modular-, or non-denitrifiers based on genes present (**Fig. 2B**). A MAG was designated complete if genes for each step of the denitrification pathway are present. A MAG was called truncated if the genome lacked only *nosZ*, suggesting the end product of nitrate metabolism is N_2_O rather than N_2_. A MAG was considered modular if it lacked *nosZ* in addition to any other genes from the denitrification pathway, suggesting it can only participate in certain reductive steps of the process. Finally, a MAG was called non-denitrifying if it contains no genes for reductive N metabolism.

### Genomic potential for complete denitrification in thermospores

Potential for complete denitrification was found in five thermospore MAGs (**Fig. 2B**). Each of the five MAGs contain a membrane-associated *nosZ* with Sec-type signal peptide characteristic of clade II enzymes [20] and all enzymes in the pathway were predicted to be located in the cytoplasmic membrane, as expected for gram-positive denitrifiers [65]. Complete denitrifiers are taxonomically classified as *Calidifontibacillus erzurumensis*, *Geobacillus thermodenitrificans*, *Neobacillus thermocopriae* and novel members of the family *Brevibacillaceae* (×2). The two *Brevibacillaceae* MAGs shared just 71.4% amino acid identity (AAI) with each other and comparison to members of this family in GTDB revealed no close relatives (>70% AAI). The greatest AAI was shared with an uncharacterised thermophilic soil bacterium, *Brevibacillaceae* species CFH-S0501 sp011059135, at 68.4 and 70.4% AAI, respectively.

Three nitric oxide reductase genes (*qnor*, *bnor*, and *snor*) were present in thermospore MAGs. The genes *bnor* and *snor* were found in genomes of complete denitrifiers in addition to *qnor* (**Fig. 2B**). Two complete denitrifiers, *Calidifontibacillus erzurumensis* and *Brevibacillaeace* (MAG 2), also contained two nitrite reductases with a *nirK* gene in addition to a *nirS* gene.

Clade II denitrifiers have a Nos gene cluster that differs to clade I denitrifiers, featuring some conserved genes that are absent in clade I genomes [21, 64]. Assessment of the Nos gene cluster (**Fig. 2C**) showed a cytochrome *c* preceding *nosZ* in all five thermospore genomes, as found in other clade II microoorganisms [46]. A gene with four transmembrane helices characteristic of *nosB* was adjacent to the putative ABC transporter complex *nosD, -Y, -F* in all of those genomes. *Calidifontibacillus erzumensis* had two Nos gene clusters, one of which had the copper chaperone *nosL* but this gene was absent from the genomes of other thermospores. The Nos gene cluster of *Brevibacillaceae* also differed from the other thermospores in that it contained a nitric oxide reductase (qNOR) immediately adjacent to the Nos gene cluster, whereas this gene was found elsewhere in the genome in the other three thermospore genomes.

### Truncated or modular denitrification potential in thermospores

Genes from the denitrification pathway were detected in five *nosZ*-lacking thermospore MAGs. These thermospores were designated truncated or modular in their metabolic potential for denitrification (**Fig. 2B**). *Symbiobacterium terraclitae* was the only MAG designated as truncated and was the only MAG to contain both *nar* and *nap* nitrate reductases (**Fig. 2B**). The *nap* nitrate reductase in *Symbiobacterium terraclitae* is monomeric (*napA*) and is distinct from the heterodimeric *napAB* commonly found in Gram-negative bacteria [66]. Ammonium-producing nitrite reductase (*nrfAH*) was present in this MAG suggesting *Symbiobacterium terraclitae* can also perform DNRA. Other MAGs designated modular have in common a respiratory nitrate reductase (membrane-bound *nar*), quinol-dependent nitric oxide reduction (*qnor*) and/or Cu-type nitrite reductase (*nirK*).

### Non-denitrifying thermospores

Two thermospore MAGs from the denitrifying enrichments contain no genes for respiratory nitrate metabolism. *Bacillus BN tepidiphilus* reached >10% relative abundance within one day of incubation and JAGHKQ01 (family *Bacillaceae G*) maintained a comparatively lower abundance (<2.5%) in all enrichments (**Fig. 2A**). Both of these genomes encode potential for glucose metabolism (mixed acid fermentation, sugar transport) and potentially became enriched by fermentative growth. Populations that ferment sugars likely provided substrates to nitrate-reducing populations in the form of fermentation products such as lactate, acetate and formate that were observed to increase in the early hours of 50°C incubations (**Fig. 1D–F**).

### Denitrification genotypes of *Bacillota*

Representative genomes from *Bacillota* phyla (*Bacillota* and *Bacillota* A−H) were retrieved from GTDB and screened for nitric oxide (cNOR, qNOR, bNOR, sNOR) and nitrous oxide reductases (NosZI, NosZII). Nitric oxide and/or nitrous oxide reductase genes were present in ∼10% of *Bacillota* genomes (*n* = 392/4216), ∼5% of *Bacillota C* genomes (*n* = 20/395), ∼4% of *Bacillota B* genomes (n = 12/323), and ∼1.5% of *Bacillota E* genomes (*n* = 5/65). Just four genomes from *Bacillota A* (*n* = 2/8243) and *Bacillota G* (*n* = 131) contained either gene and all genomes from the phyla *Bacillota D*, *F*, and *H* (170 genomes) lacked both.

Genomes with nitric oxide and/or nitrous oxide reductases were screened with the full denitrification HMM set and included in a phylogenomic tree with MAGs from this study (**Fig. 3**). Genomic potential for complete denitrification is constrained to the phylum *Bacillota* and was only found within members of the class *Bacilli*. Seventeen genera contain complete denitrifiers (**Fig. 3**; 41 GTDB MAGs + 5 MAGs from this study). This includes 11 genomes that encode bNOR and/or sNOR but lack qNOR (or cNOR) and would have been considered incomplete denitrifiers using KO annotations only. Cytochrome *c*-dependent nitric oxide reductase (cNOR) was only encoded in genomes within the family *Desulfitobacteriaceae* from the phylum *Bacillota B* (**Table S2**). *Desulfitobacteriaceae* also contain clade II-type *nosZ* but lack genes for nitrite reduction.

**Figure 3:**
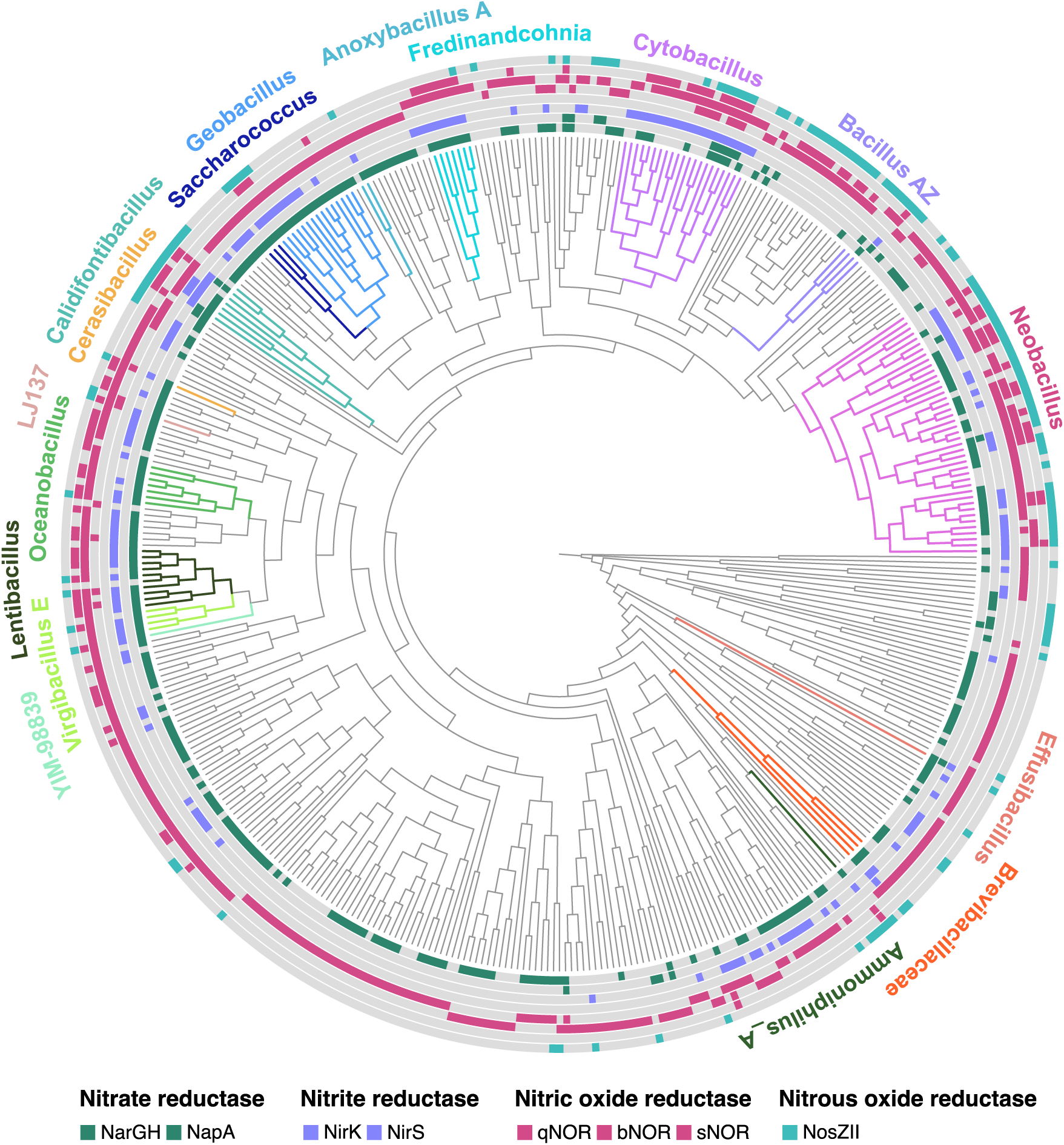
*Bacillota* genera with genomic potential for complete denitrification. A phylogenomic tree was constructed with *Bacillota* MAGs from this study and 433 genomes from GTDB that contain nitric oxide reductase and/or nitrous oxide reductase. Coloured wedges at the tips of branches indicate gene presence (filled) or absence (grey). Genes shown from the innermost circle to the outer: *narGH*, *napA*, *nirK*, *nirS*, *qnor, bnor, snor,* clade II-type *nosZ*. The phylogenomic tree was constructed using a concatenated alignment of 119 single copy genes conserved within *Bacillota* and is presented as a cladogram. Within the phylum *Bacillota*, 17 genera have potential for complete denitrification (coloured clades with bold text).

Among denitrification genotypes, the absence of one or more of the reduction steps is common. A further 74 MAGs were categorised as truncated denitrifiers i.e., missing only nitrous oxide reductase, 73 were categorised as nitric oxide reducers i.e., contain nitric oxide reductase only, and 15 were categorised as non-denitrifying nitrous oxide reducers i.e., contain nitrous oxide reductase only (**Fig. 3 and Table S5**). The remaining genomes were categorised as modular i.e., they contain nitric oxide reductase and/or nitrous oxide reductase in addition to one or more denitrification pathway genes. Many *Bacillota* genomes encode qNOR (172/433) or bNOR (123/433) whereas sNOR was rarely found without the presence of another nitric oxide reductase gene (2/433). Occurrences of three NOR genes in the same genome (qNOR, bNOR and sNOR) was observed in 26 genomes, most of which belong to *Neobacillus* and other genera containing complete denitrifiers (**Fig. 3**).

## Discussion

Targeted enrichment with nitrate resulted in the germination and activity of denitrifying thermospore populations. This approach uncovered multiple lineages of denitrifiers, including novel members of the family *Brevibacillaceae.* Different thermospore taxa responded in parallel incubations suggesting there are heterogeneous populations of dormant thermospores in Athabasca oil sands outcrops, which is consistent with similar observations of oil sands microbiomes generally [25, 67]. It is well documented that thermospores comprise part of the microbial seedbank in cold sedimentary [68, 69] and soil environments [70, 71]. When conditions change favourably, dormant populations germinate and become active members of the microbial community. This premise underpins strategies for engineered microbial activity *in situ* with the objective of pressure generation and maintenance via microbial biogas such as N_2_ [72]. Our results demonstrate the feasibility for denitrifying thermospores to be stimulated in oil sands.

Denitrifying thermospores enriched from oil sands have clade II-type *nosZ* genes (subclade H) for catalysing reduction of N_2_O to N_2_. Clade II *nosZ*-bearing microorganisms are numerically significant in the environment [73–76], though they are often considered to be non-denitrifiers lacking genes needed for the stepwise reduction of nitrate to dinitrogen [21, 77]. While this is true for certain lineages within the diverse clade II-type NosZ, analysis of MAGs from N_2_-producing enrichments in this study, as well as *Bacillota* from GTDB, shows that multiple genera within the class *Bacilli* contain a full complement of denitrification genes. This is consistent with multiple isolated representatives from this class that have been experimentally shown to perform complete denitrification [22]. Examples include *Geobacillus thermodenitrificans* isolated from a deep oil reservoir [78] and *Calidifontibacillus azotoformans* (formerly *Bacillus azotoformans*) isolated from soil [65].

Genomic potential for complete denitrification was present in 17 genera within the class *Bacilli*. Within these genera, it was common for microorganisms to possess multiple nitric oxide reductases (qNOR, bNOR and sNOR). This highlights that NOR enzymes are not mutually exclusive, though the different conditions under which they are not expressed in *Bacillota* are not clear. It has been suggested that bNOR in *Bacillus azotoformans* can be used for aerobic NO reduction in microoxic environments [46]. The recently characterised enzyme eNOR, that descends from the same family of oxygen reductases as bNOR, also reduces nitric oxide under aerobic conditions [19, 79]. Members of the *Bacillota* are often identified as contributors to denitrification in environments with variable redox conditions, including agricultural soil, deep vadose zone soil, and rice paddy soil [7, 80, 81] and are widespread in the soil microbiome generally [82]. Having multiple nitric oxide reductases could provide *Bacillota* with metabolic versatility in environments like soils, where combined oxic and anoxic conditions are commonly found [3].

So-called functional redundancy has also been found with other enzymes within the denitrification pathway. *B. azotoformans* contains five Nos gene clusters, three of which include a *nosZ* gene (Heylen 2012). In addition, a recent survey of nitrite reductases (NirK and NirS) in isolates and MAGs showed that possessing both enzymes is more common than previously appreciated and potentially allows microorganisms bearing both enzymes to denitrify across a wider range of environments [83]. Functionally redundant enzymes within a genome may also reflect an ability of the microorganisms to adapt to changing environmental conditions. This would be a beneficial trait for members of the *Bacillota* as endospore-formers undergo periods of dormancy and respond rapidly through germination to changes in their environment.

Genome-resolved metagenomics is a useful approach for studying denitrification as it provides the gene content of populations and circumvents challenges with PCR-based marker gene approaches. However, we found that certain denitrification genes were missed, or pathways appeared incomplete, using standard annotation databases that are biased towards clade I denitrifiers. For example, the denitrification reference pathway in KEGG includes nitrate reductase composed of subunits *napAH* and nitric oxide reductase composed of subunits *norBC*. However, nitrate reductases in *Symbiobacterium* (*Bacillota E*) are monomeric [66] and lack the *napH* subunit. Similarly, qNOR nitric oxide reductases are fused and lack the *norC* subunit [15]. This can result in modules or pathways appearing incomplete. The bNOR nitric oxide reductase has to date only been found in *Bacillota* and was originally isolated from *Bacillus azotoformans* [18, 65]. Despite being biochemically characterised bNOR genes were not identified with commonly used gene annotation databases KEGG, eggNOG or TIGRfam. Finally, while putative NOR enzyme families that have been recently proposed [19, 79] are not expected to be represented in curated annotation databases, their absence nevertheless highlights that interpretation of community gene content is limited by the breadth of gene databases. Considering the complete diversity of NOR enzymes reveals a greater diversity of microorganisms capable of denitrification. This is an important consideration for studies attempting to quantify denitrification or N_2_O emissions to ensure that metabolic potential is not underestimated.

## Acknowledgements

This work was funded by a Campus Alberta Innovates Program (CAIP) chair awarded to CRJH.

## Data Availability Statement

DNA sequencing data (16S rRNA gene amplicon, metagenome, and metagenome-assembled genomes) are available at the NCBI Sequence Read Archive (SRA) under BioProject ID PRJNA1110647.

## Notes

### Competing Interest Statement

CRJH and MF have granted patents for an oil recovery technology that depends on biogas production by thermophilic bacteria, related to the presented research. Patents are owned by a start-up company in which CRJH and MF are co-founders. EB and JC declare no potential competing interests.

## References

1. Tian H, Xu R, Canadell JG, Thompson RL, Winiwarter W, Suntharalingam P, et al. A comprehensive quantification of global nitrous oxide sources and sinks. Nature 2020; 586: 248– 256.

2. Harris E, Yu L, Wang Y-P, Mohn J, Henne S, Bai E, et al. Warming and redistribution of nitrogen inputs drive an increase in terrestrial nitrous oxide emission factor. Nat Commun 2022; 13: 4310.

3. Christensen S, Rousk K. Global N2O emissions from our planet: Which fluxes are affected by man, and can we reduce these? iScience 2024; 27.

4. Bowles TM, Atallah SS, Campbell EE, Gaudin ACM, Wieder WR, Grandy AS. Addressing agricultural nitrogen losses in a changing climate. Nat Sustain 2018; 1: 399–408.

5. Nilsson JE, Weisner SEB, Liess A. Wetland nitrogen removal from agricultural runoff in a changing climate. Sci Total Environ 2023; 892: 164336.

6. Conthe M, Lycus P, Arntzen MØ, Ramos da Silva A, Frostegård Å, Bakken LR, et al. Denitrification as an N2O sink. Water Res 2019; 151: 381–387.

7. Zhang L, Zhao H, Qin S, Hu C, Shen Y, Qu B, et al. Genome-Resolved Metagenomics and Denitrifying Strain Isolation Reveal New Insights into Microbial Denitrification in the Deep Vadose Zone. Environ Sci Technol 2024.

8. Philippot L. Denitrifying genes in bacterial and Archaeal genomes. Biochim Biophys Acta BBA - Gene Struct Expr 2002; 1577: 355–376.

9. Chee-Sanford JC, Connor L, Krichels A, Yang WH, Sanford RA. Hierarchical detection of diverse Clade II (atypical) nosZ genes using new primer sets for classical- and multiplex PCR array applications. J Microbiol Methods 2020; 172: 105908.

10. Heylen K, Gevers D, Vanparys B, Wittebolle L, Geets J, Boon N, et al. The incidence of nirS and nirK and their genetic heterogeneity in cultivated denitrifiers. Environ Microbiol 2006; 8: 2012– 2021.

11. Wei W, Isobe K, Nishizawa T, Zhu L, Shiratori Y, Ohte N, et al. Higher diversity and abundance of denitrifying microorganisms in environments than considered previously. ISME J 2015; 9: 1954–1965.

12. Decleyre H, Heylen K, Tytgat B, Willems A. Highly diverse nirK genes comprise two major clades that harbour ammonium-producing denitrifiers. BMC Genomics 2016; 17: 527.

13. Sun H, Jiang S. A review on nirS-type and nirK-type denitrifiers via a scientometric approach coupled with case studies. Environ Sci Process Impacts 2022; 24: 221–232.

14. Saraste M, Castresana J. Cytochrome oxidase evolved by tinkering with denitrification enzymes. FEBS Lett 1994; 341: 1–4.

15. Hemp J, Gennis RB. Diversity of the Heme–Copper Superfamily in Archaea: Insights from Genomicsand Structural Modeling. In: Schäfer G, Penefsky HS (eds). Bioenergetics: Energy Conservation and Conversion. 2008. Springer Berlin Heidelberg, Berlin, Heidelberg, pp 1–31.

16. Hino T, Matsumoto Y, Nagano S, Sugimoto H, Fukumori Y, Murata T, et al. Structural Basis of Biological N2O Generation by Bacterial Nitric Oxide Reductase. Science 2010; 330: 1666–1670.

17. Matsumoto Y, Tosha T, Pisliakov AV, Hino T, Sugimoto H, Nagano S, et al. Crystal structure of quinol-dependent nitric oxide reductase from Geobacillus stearothermophilus. Nat Struct Mol Biol 2012; 19: 238–245.

18. Al-Attar S, de Vries S. An electrogenic nitric oxide reductase. FEBS Lett 2015; 589: 2050–2057.

19. Murali R, Pace LA, Sanford RA, Ward LM, Lynes M, Hatzenpichler R, et al. Diversity and evolution of nitric oxide reduction. bioRxiv 2021; 2021.10.15.464467.

20. Sanford RA, Wagner DD, Wu Q, Chee-Sanford JC, Thomas SH, Cruz-García C, et al. Unexpected nondenitrifier nitrous oxide reductase gene diversity and abundance in soils. Proc Natl Acad Sci 2012; 109: 19709 LP – 19714.

21. Hallin S, Philippot L, Löffler FE, Sanford RA, Jones CM. Genomics and Ecology of Novel N2O-Reducing Microorganisms. Trends Microbiol 2018; 26: 43–55.

22. Verbaendert I, Hoefman S, Boeckx P, Boon N, De Vos P. Primers for overlooked nirK, qnorB, and nosZ genes of thermophilic Gram-positive denitrifiers. FEMS Microbiol Ecol 2014; 89: 162–180.

23. Ma Y, Zilles JL, Kent AD. An evaluation of primers for detecting denitrifiers via their functional genes. Environ Microbiol 2019; 21: 1196–1210.

24. Wong ML, An D, Caffrey SM, Soh J, Dong X, Sensen CW, et al. Roles of thermophiles and fungi in bitumen degradation in mostly cold oil sands outcrops. Appl Environ Microbiol 2015; 81: 6825– 6838.

25. Ridley CM, Voordouw G. Aerobic microbial taxa dominate deep subsurface cores from the Alberta oil sands. FEMS Microbiol Ecol 2018; 94.

26. Caporaso JG, Lauber CL, Walters WA, Berg-Lyons D, Lozupone CA, Turnbaugh PJ, et al. Global patterns of 16S rRNA diversity at a depth of millions of sequences per sample. Proc Natl Acad Sci 2011; 108: 4516–4522.

27. Martin M. Cutadapt removes adapter sequences from high-throughput sequencing reads. EMBnet.journal 2011; 17: 10.

28. Callahan BJ, McMurdie PJ, Rosen MJ, Han AW, Johnson AJA, Holmes SP. DADA2: High-resolution sample inference from Illumina amplicon data. Nat Methods 2016; 13: 581–583.

29. Krueger F, James F, Ewels P, Afyounian E, Weinstein M, Schuster-Boeckler B, et al. FelixKrueger/TrimGalore: v0.6.10 - add default decompression path. 2023. Zenodo.

30. Li D, Liu CM, Luo R, Sadakane K, Lam TW. MEGAHIT: An ultra-fast single-node solution for large and complex metagenomics assembly via succinct de Bruijn graph. Bioinformatics 2015; 31: 1674–1676.

31. Bushnell B, Rood J, Singer E. BBTools Software Package. Plos One. https://sourceforge.net/projects/bbmap/.

32. Kang DD, Li F, Kirton E, Thomas A, Egan R, An H, et al. MetaBAT 2: An adaptive binning algorithm for robust and efficient genome reconstruction from metagenome assemblies. PeerJ 2019; 2019: e7359–e7359.

33. Alneberg J, Bjarnason BS, De Bruijn I, Schirmer M, Quick J, Ijaz UZ, et al. Binning metagenomic contigs by coverage and composition. Nat Methods 2014; 11: 1144–1146.

34. Sieber CMKK, Probst AJ, Sharrar A, Thomas BC, Hess M, Tringe SG, et al. Recovery of genomes from metagenomes via a dereplication, aggregation and scoring strategy. Nat Microbiol 2018; 3: 836–843.

35. Chklovski A, Parks DH, Woodcroft BJ, Tyson GW. CheckM2: a rapid, scalable and accurate tool for assessing microbial genome quality using machine learning. bioRxiv 2022; 2022.07.11.499243.

36. Olm MR, Brown CT, Brooks B, Banfield JF. dRep: a tool for fast and accurate genomic comparisons that enables improved genome recovery from metagenomes through de-replication. ISME J 2017; 11: 2864–2868.

37. Aroney STN, Newell RJP, Nissen J, Camargo AP, Tyson GW, Woodcroft BJ. CoverM: Read coverage calculator for metagenomics. 2024. Zenodo.

38. Chaumeil P-A, Mussig AJ, Hugenholtz P, Parks DH. GTDB-Tk v2: memory friendly classification with the genome taxonomy database. Bioinformatics 2022; 38: 5315–5316.

39. Rodriguez-R LM, Konstantinidis KT. The enveomics collection: a toolbox for specialized analyses of microbial genomes and metagenomes. PeerJ Prepr 2016; 4: e1900v1.

40. Kanehisa M, Sato Y, Morishima K. BlastKOALA and GhostKOALA: KEGG Tools for Functional Characterization of Genome and Metagenome Sequences. J Mol Biol 2016; 428: 726–731.

41. Huerta-Cepas J, Szklarczyk D, Heller D, Hernández-Plaza A, Forslund SK, Cook H, et al. eggNOG 5.0: a hierarchical, functionally and phylogenetically annotated orthology resource based on 5090 organisms and 2502 viruses. Nucleic Acids Res 2019; 47: D309–D314.

42. Li G, Rabe KS, Nielsen J, Engqvist MKM. Machine Learning Applied to Predicting Microorganism Growth Temperatures and Enzyme Catalytic Optima. ACS Synth Biol 2019; 8: 1411–1420.

43. Teufel F, Almagro Armenteros JJ, Johansen AR, Gíslason MH, Pihl SI, Tsirigos KD, et al. SignalP 6.0 predicts all five types of signal peptides using protein language models. Nat Biotechnol 2022; 40: 1023–1025.

44. Hallgren J, Tsirigos KD, Pedersen MD, Armenteros JJA, Marcatili P, Nielsen H, et al. DeepTMHMM predicts alpha and beta transmembrane proteins using deep neural networks. 2022. bioRxiv., 2022.04.08.487609

45. Hein S, Witt S, Simon J. Clade II nitrous oxide respiration of Wolinella succinogenes depends on the NosG, -C1, -C2, -H electron transport module, NosB and a Rieske/cytochrome bc complex. Environ Microbiol 2017; 19: 4913–4925.

46. Heylen K, Keltjens J. Redundancy and modularity in membrane-associated dissimilatory nitrate reduction in Bacillus. Front Microbiol 2012; 3: 371.

47. Yu NY, Wagner JR, Laird MR, Melli G, Rey S, Lo R, et al. PSORTb 3.0: improved protein subcellular localization prediction with refined localization subcategories and predictive capabilities for all prokaryotes. Bioinforma Oxf Engl 2010; 26: 1608–1615.

48. Wilkins D. gggenes: Draw Gene Arrow Maps in ‘ggplot2’. 2023.

49. Wickham H. ggplot2: Elegant Graphics for Data Analysis. 2016.

50. Rinaldo S, Giardina G, Castiglione N, Stelitano V, Cutruzzolà F. The catalytic mechanism of Pseudomonas aeruginosa cd1 nitrite reductase. Biochem Soc Trans 2011; 39: 195–200.

51. Gomaa F, Utter DR, Powers C, Beaudoin DJ, Edgcomb VP, Filipsson HL, et al. Multiple integrated metabolic strategies allow foraminiferan protists to thrive in anoxic marine sediments. Sci Adv 2021; 7: eabf1586.

52. Sievers F, Wilm A, Dineen D, Gibson TJ, Karplus K, Li W, et al. Fast, scalable generation of high-quality protein multiple sequence alignments using Clustal Omega. Mol Syst Biol 2011; 7: 539.

53. Larsson A. AliView: a fast and lightweight alignment viewer and editor for large datasets. Bioinformatics 2014; 30: 3276–3278.

54. Eddy SR. Accelerated Profile HMM Searches. PLoS Comput Biol 2011; 7: e1002195–e1002195.

55. Parks DH, Chuvochina M, Rinke C, Mussig AJ, Chaumeil P-A, Hugenholtz P. GTDB: an ongoing census of bacterial and archaeal diversity through a phylogenetically consistent, rank normalized and complete genome-based taxonomy. Nucleic Acids Res 2022; 50: D785–D794.

56. Lee MD. GToTree: A user-friendly workflow for phylogenomics. Bioinformatics 2019; 35: 4162–4164.

57. Edgar RC. Search and clustering orders of magnitude faster than BLAST. Bioinformatics 2010; 26: 2460–2461.

58. Selengut JD, Haft DH, Davidsen T, Ganapathy A, Gwinn-Giglio M, Nelson WC, et al. TIGRFAMs and Genome Properties: tools for the assignment of molecular function and biological process in prokaryotic genomes. Nucleic Acids Res 2007; 35: D260–D264.

59. Hyatt D, Chen G-L, Locascio PF, Land ML, Larimer FW, Hauser LJ. Prodigal: prokaryotic gene recognition and translation initiation site identification. BMC Bioinformatics 2010; 11: 119.

60. Robert C. Edgar. High-accuracy alignment ensembles enable unbiased assessments of sequence homology and phylogeny. bioRxiv 2022; 2021.06.20.449169.

61. Capella-Gutiérrez S, Silla-Martínez JM, Gabaldón T. trimAl: a tool for automated alignment trimming in large-scale phylogenetic analyses. Bioinformatics 2009; 25: 1972–1973.

62. Price MN, Dehal PS, Arkin AP. FastTree 2 – Approximately Maximum-Likelihood Trees for Large Alignments. PLOS ONE 2010; 5: e9490.

63. Bianchini G, Sánchez-Baracaldo P. TreeViewer: Flexible, modular software to visualise and manipulate phylogenetic trees. Ecol Evol 2024; 14: e10873.

64. Hein S, Simon J. Chapter Four - Bacterial nitrous oxide respiration: electron transport chains and copper transfer reactions. In: Poole RK (ed). Advances in Microbial Physiology. 2019. Academic Press, pp 137–175.

65. Suharti, de Vries S. Membrane-bound denitrification in the Gram-positive bacterium Bacillus azotoformans. Biochem Soc Trans 2005; 33: 130–133.

66. Jepson BJN, Marietou A, Mohan S, Cole JA, Butler CS, Richardson DJ. Evolution of the soluble nitrate reductase: defining the monomeric periplasmic nitrate reductase subgroup. Biochem Soc Trans 2006; 34: 122–126.

67. An D, Caffrey SM, Soh J, Agrawal A, Brown D, Budwill K, et al. Metagenomics of Hydrocarbon Resource Environments Indicates Aerobic Taxa and Genes to be Unexpectedly Common. 2013.

68. Hubert C, Loy A, Nickel M, Arnosti C, Baranyi C, Bruchert V, et al. A Constant Flux of Diverse Thermophilic Bacteria into the Cold Arctic Seabed. Science 2009; 325: 1541–1544.

69. Müller AL, de Rezende JR, Hubert CRJ, Kjeldsen KU, Lagkouvardos I, Berry D, et al. Endospores of thermophilic bacteria as tracers of microbial dispersal by ocean currents. ISME J 2014; 8: 1153– 1165.

70. Marchant R, Banat IM, Rahman TJ, Berzano M. The frequency and characteristics of highly thermophilic bacteria in cool soil environments. Environ Microbiol 2002; 4: 595–602.

71. Marchant R, Franzetti A, Pavlostathis SG, Tas DO, Erdbrugger I, Unyayar A, et al. Thermophilic bacteria in cool temperate soils: Are they metabolically active or continually added by global atmospheric transport? Appl Microbiol Biotechnol 2008; 78: 841–852.

72. Hubert CRJ, Fustic M. Microbially enhanced thermal oil recovery. 2021. Washington, DC.

73. Jones CM, Graf DR, Bru D, Philippot L, Hallin S. The unaccounted yet abundant nitrous oxide-reducing microbial community: a potential nitrous oxide sink. ISME J 2013; 7: 417–426.

74. Coyotzi S, Doxey AC, Clark ID, Lapen DR, Van Cappellen P, Neufeld JD. Agricultural soil denitrifiers possess extensive nitrite reductase gene diversity. Environ Microbiol 2017; 19: 1189– 1208.

75. Mosley OE, Gios E, Close M, Weaver L, Daughney C, Handley KM. Nitrogen cycling and microbial cooperation in the terrestrial subsurface. ISME J 2022; 1–13.

76. Tang W, Jayakumar A, Sun X, Tracey JC, Carroll J, Wallace E, et al. Nitrous Oxide Consumption in Oxygenated and Anoxic Estuarine Waters. Geophys Res Lett 2022; 49: e2022GL100657.

77. Conthe M, Wittorf L, Kuenen JG, Kleerebezem R, van Loosdrecht MCM, Hallin S. Life on N2O: deciphering the ecophysiology of N2O respiring bacterial communities in a continuous culture. ISME J 2018; 12: 1142–1153.

78. Feng L, Wang W, Cheng J, Ren Y, Zhao G, Gao C, et al. Genome and proteome of long-chain alkane degrading Geobacillus thermodenitrificans NG80-2 isolated from a deep-subsurface oil reservoir. Proc Natl Acad Sci U S A 2007; 104: 5602–5607.

79. Murali R, Hemp J, Gennis RB. Evolution of quinol oxidation within the heme-copper oxidoreductase superfamily. Biochim Biophys Acta BBA - Bioenerg 2022; 1863: 148907.

80. Ishii Satoshi, Yamamoto Michihiro, Kikuchi Mami, Oshima Kenshiro, Hattori Masahira, Otsuka Shigeto, et al. Microbial Populations Responsive to Denitrification-Inducing Conditions in Rice Paddy Soil, as Revealed by Comparative 16S rRNA Gene Analysis. Appl Environ Microbiol 2009; 75: 7070–7078.

81. Anderson CR, Peterson ME, Frampton RA, Bulman SR, Keenan S, Curtin D. Rapid increases in soil pH solubilise organic matter, dramatically increase denitrification potential and strongly stimulate microorganisms from the Firmicutes phylum. PeerJ 2018; 6: e6090.

82. Bandopadhyay S, Shade A. Chapter 3 - Soil bacteria and archaea. In: Paul EA, Frey SD (eds). Soil Microbiology, Ecology and Biochemistry (Fifth Edition). 2024. Elsevier, pp 41–74.

83. Pold G, Bonilla-Rosso G, Saghaï A, Strous M, Jones CM, Hallin S. Phylogenetics and environmental distribution of nitric oxide forming nitrite reductases reveals their distinct functional and ecological roles. ISME Commun 2024; ycae020.

